# iSleep: Continuous, binocular pupil tracking in sleep and reduced consciousness for physiological monitoring, predictions and interventions

**DOI:** 10.1101/2025.06.13.659058

**Authors:** Ozge Yuzgec, Guillaume Legendre, Amirhossein Asadian, Yesica Gloria, Laurence Bayer, Quentin Tourdot, Sebastien Pellat, Sahsa Novozhilova, Camille D’Ancona, Marina Ulanova, Mateusz Kecik, Andrea Maccari, David Lopez, Laura Licini, Gaël Nguyen Tang, Sébastien Van Delden, Giorgio Enrico Bravetti, Tin Vo, Martina Kropp, Colette Boex, Line Jakus, Karl Schaller, Fanny Sollander, Laszlo Vutskits, Gabriele Thumann, Sophie Schwartz, Daniel Huber

## Abstract

Monitoring pupil dynamics is a key tool in understanding arousal. Pupil size can serve as a biomarker for the autonomic nervous system balance as well as for identifying brain states. While internal states can also be self-reported when awake, automated detection and non-invasive monitoring is crucial during sleep and reduced consciousness. Here, we introduce iSleep, an innovative pupil tracking and analysis framework for sleep in humans. It features comfortable, humidified eye-tracking goggles and a platform for integrated analysis and prediction capabilities. We show that iSleep allows safe and continuous access to binocular pupil size and ocular dynamics during sleep and anesthesia. iSleep reveals that pupillary fluctuations correlate tightly with brain activity, heartbeat, and breathing, and can reliably predict brain states. Pupil constrictions reflect parasympathetic drive and likely serve a protective function for deep sleep stability; while dilations indicate arousals. Unexpectedly, we observed a decoupling of binocular movements during periods of sleep, indicating alterations in reflexes which usually govern voluntary eye movements. Finally, iSleep was tested in surgery patients under general anesthesia, revealing dynamic pupil changes to noxious stimuli, suggesting the potential for nociception monitoring during surgeries. In summary, iSleep offers an easy-to-use, robust alternative to read out brain states during sleep and anesthesia, opening new avenues in monitoring, diagnostics, and treatments, previously obscured by closed eyelids.

## Introduction

Pupil size has long been used as a physiological marker for attention, arousal, emotion or cognitive performance [1]–[3]. It not only tightly correlates with EEG activity (i.e. brain states) across various regions, it also co-fluctuates with measures of the autonomic nervous system such as heart rate (HR) and skin conductance [1], [4]–[6]. Combining neurophysiology and imaging techniques in animal models further revealed that pupil constrictions reflect cholinergic activity, whereas dilations are timed to increased locus coeruleus activity or locomotion [7]–[10]. In parallel, eye tracking research provides ample evidence that eye movements may be strong indicators of cognitive processing or disease states [11]–[13]. Together, these findings led to the use of pupil dynamics as predictive measures for behavior, physiology and continuous markers of brain and autonomic states in research [8], [10], [14]–[17] and commercial settings [18], [19].

While pupil size is a reliable biomarker in the awake state, its use during sleep or unconsciousness has been limited by eyelid occlusion. Although pupils are regularly examined by manual opening in clinical settings such as in emergency rooms, operating theaters or intensive care units [20], a systematic and continuous readout of eye kinematics is currently not available. Thus, clinicians and researchers rely heavily on other physiological recordings during sleep and under general anesthesia. In particular, the current gold standard in monitoring remains polysomnography (PSG), which combines electroencephalography (EEG), electro-oculography (EOG), and electromyography (EMG) [21], [22]. This technique involves a relatively complex, albeit standardized system of electrodes, cables, amplifier filters and analysis systems. Alternative methods including biosensors such as skin conductance electrodes [23] or reduced EEGs like the bispectral index system (BIS) are also regularly used [24]. Compared to these electrophysiological systems, pupil size measurements can serve as a complementary biomarker during interventions and provide an additional layer of insight, enhancing the overall efficacy and precision of patient assessments in various clinical situations.

We therefore asked if it is possible to track pupil dynamics during sleep in humans and use them as biomarkers to determine brain states. Here, we describe iSleep, the first continuous, binocular pupil tracking system for sleeping humans. It combines binocular eye tracking hardware, an extensive pupil-PSG database and an analysis framework, which allows automated detection of brain states based on pupil parameters during sleep utilizing the iSleep database. We illustrate a series of applications and provide important biological insights using this system. We show (i) that iSleep is a practical, safe and comfortable system, capable of providing reliable measurements of spontaneous and sensory evoked pupillary changes in sleep, (ii) that these dynamics can be used as a reliable proxy to read out, and (iii) predict brain states and the activity of the autonomic nervous system thus providing novel biomarkers for altered states of consciousness. Using iSleep’s combined visual stimulation and monitoring capacities, we also demonstrate that (iv) the light-evoked pupil constriction persists during sleep and relates to stabilizing sleep states. In addition, gaze tracking with iSleep allowed us to reveal (v) disconjugate eye movements during large parts of NREM and REM sleep. Finally, when using iSleep during surgery under general anesthesia, we could detect (vi) pupil responses (dilations) to noxious stimuli. Taken together, with its high-accuracy prediction performance and real-time monitoring capabilities, the iSleep system represents a powerful research tool for sleep and consciousness studies. It furthermore offers novel biomarkers to assist clinicians during interventions.

## Results

### iSleep goggles allow safe open-eye pupil recordings during sleep

Inspired by our previous work on pupil tracking for sleeping mice [10], we developed a novel optical approach able to record binocular eye movements and pupil dynamics during sleep in humans [25]. The system, we named iSleep (Fig. 1), consists of form-fitted goggles containing two miniature cameras and infrared lights to illuminate the eyes (Fig. 1a & b). To gain optical access to the eyes, the eyelids were held open with soft, elastic medical tape (Fig. 1c), which permitted the subjects to blink when necessary. The enclosed environment of the iSleep goggles was humidified by inserting single-use wet sponges into side-mounted holders. Temperature (34-36°C) and humidity levels (80-95%) were continuously monitored using electronic sensors (Fig. 1d). This enabled prolonged and safe recordings of open eye data, including pupil size and eye position (Fig. 1e). We first assessed if sleeping with the iSleep goggles and with open eyelids would be possible and if so, whether it affected the overall sleep patterns. We found that participants exhibited comparable sleep architecture in open-eye and closed-eye naps (Fig. S1a). Polysomnographic activity displayed usual characteristics of each sleep stage, regarding variations in EEG activity, muscle tone, eye movements, breathing and cardiac activity (Fig. 1e-f). Critically, sleep architecture did not differ when comparing open and closed-eye naps (percentage time spent across different stages, student’s t-test n.s. WAKE p=0.988, NREM1 p= 0.067, NREM2 p=0.261, NREM3 p=0.189, REM p=0.342; duration of each sleep bout, student’s t-test WAKE p=0.265, NREM1 p=0.718, NREM2 p =0.709, NREM3 p=0.810, REM p=0.872; fragmentation p = 0.509, student’s t-test, Supplementary Fig. 1).

**Figure 1.**
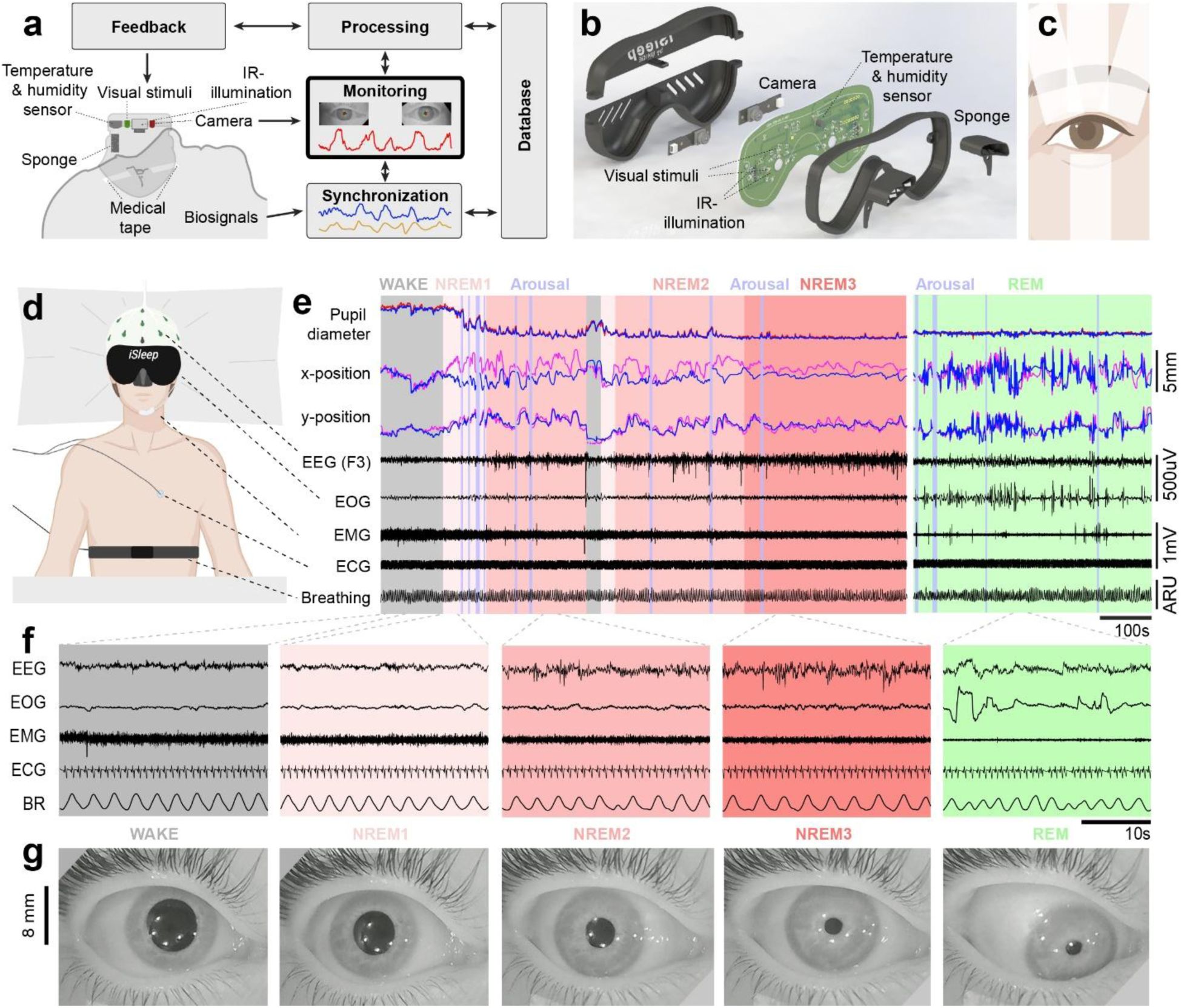
iSleep framework as a non-invasive, modular and closed-loop system for monitoring pupils during sleep and reduced consciousness. a. iSleep framework encompasses (i) a humidified, temperature controlled, binocular eye monitoring sleep goggle that can also provide visual stimulation, (ii) analytics modules for data processing and synchronization, and (iii) database that includes synchronized streams PSG signals as well as pupil tracking data. b. Exploded schematic illustration of the iSleep goggle components. c. Opening of the eyes with soft, medical tapes. d. Participant with iSleep and PSG setup. e. Pupil diameter, horizontal (x) and vertical (y) eye position, and PSG recording of the participant during a nap. f. EEG, EOG, EMG, ECG and BR signals depicting the natural characteristics of each sleep stage during wake and open-eye sleep. g. Snapshots from the video recording of the participant’s right eye across wake and sleep stages.

As a measure of tolerance and comfort, we compared participants’ subjective reports, in which they ranked the quality of their sleep and eye comfort in closed- and open-eye naps (N=17 participants, n=50 sessions). Subjective ratings of participants’ sleep quality was comparable across both types of naps (Supplementary Fig. 1d, 82% open-eye nap with good sleep, 95% closed-eye nap with good sleep, n.s. p = 0.473, binomial test). Participants also reported that they felt similarly well rested (Supplementary Fig. 1d). Whereas eye comfort and humidity was also overall in a comparable range, closed-eye sleep was ranked better (Supplementary Fig. 1d, 70% good or more, 30% “acceptable” in open-eye naps; 100% good or more in closed-eye naps). These responses confirmed that open-eye pupil tracking with iSleep did not significantly alter sleep architecture and provided comfortable conditions in a controlled research environment (Methods, Fig. 1g).

### Continuous tracking of pupil dynamics during sleep reveals high correlations with brain activity and autonomic rhythms

Previous research on mice during quiet-wakefulness and sleep, as well as on awake humans revealed a robust relationship between the pupil size fluctuations and EEG markers of arousal, whereby pupil diameter correlates negatively with low frequency power and positively with higher frequency power [7], [14], [26], [27]. Yet, whether this relationship also holds during sleep in humans is unknown. We thus combined the iSleep framework with other physiological measures including EEG, heart rate and breathing. The generation of a large database of synchronously acquired parameters allowed determining if pupil size can be used as a biomarker of brain states and other physiological rhythms.

We first performed cross-correlation analysis of pupil size with EEG activity across all sleep stages (Fig 2a-b). We found a marked negative correlation with markers of sleep depth (on average, r= −0.17 for delta, r=−0.13 for theta and r=−0.22 for sigma frequency bands, p<0.001) during NREM sleep, with the strongest negative correlation observed for the sigma band (Fig 2b, significant cluster of negative correlation from −0.4 to 20 s lag; t_cluster_=−1944.564, p<0.001). In other words, decreases in pupil diameter coincide with increases in sigma power. Weaker but similar correlation patterns were observed during WAKE (r= −0.17 with delta power, p<0.001) and REM (r= −0.06 with delta power, p=0.119) states (Fig 2b). Conversely, we found positive correlations between pupil diameter and higher frequency bands linked to increased arousal (on average, r=0.13 p=0.001, r=0.04 p<0.001, r=0.14 p<0.001 with alpha power in wakefulness, NREM and REM, respectively, and r=0.08, r=0.16, r=0.07 with beta power in wakefulness, NREM and REM, respectively) [28], [29].

**Figure 2.**
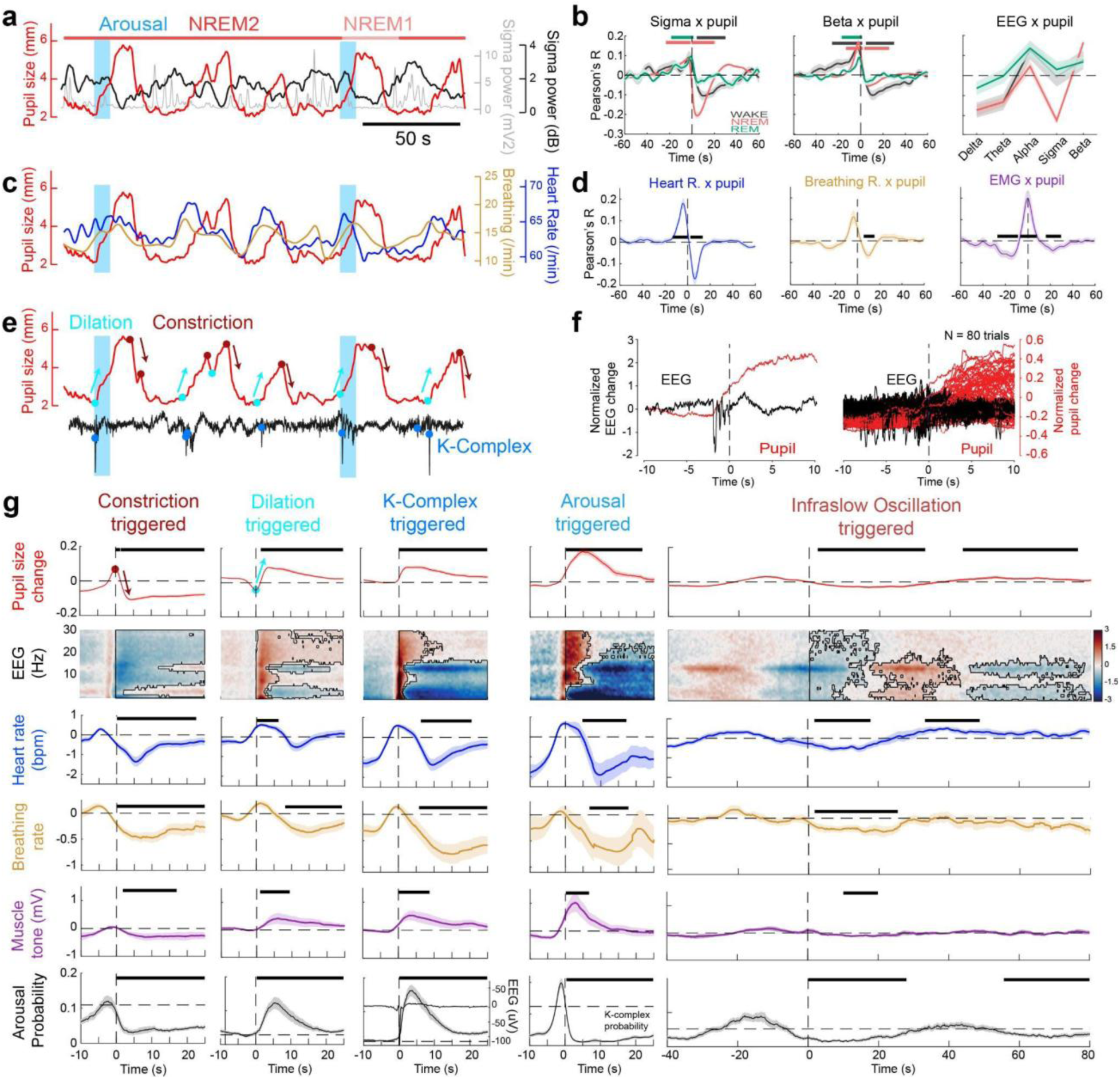
Pupil tracking during sleep reveals tight links to EEG and autonomic nervous system activity. **a.** NREM sleep bout from one participant illustrating co-fluctuations of pupil size (red), frontal EEG sigma power (gray) and its low-pass filtered trace (black). **b.** Pupil size vs. sigma power correlations across wakefulness, NREM and REM sleep (left), Pearson’s R average of selected EEG bands (middle), lags of peak correlations for pupil vs. selected EEG bands (right) (N=19 participants and n=45 sessions). **c.** Pupil size, HR and breathing rate (BR) in NREM sleep (same participant and NREM bout as in a.). **d.** Cross-correlations of HR (left, N=19 n=45), BR (middle, N=8, n=22) and EMG activity (right, N=19, n=39) with pupil size in NREM sleep. **e.** Pupil size and EEG traces in NREM illustrating rapid dilations and constrictions in relation to microarousals and K-complexes (same participant and NREM bout as in a.). **f.** Left: Single trial example of simultaneous occurrence of a K-complex and pupil dilation during an arousal. Right: the overlay of 80 arousal examples from 5 sessions from 4 participants. Quantification of speed of pupil constriction and dilations in different sleep stages. Note the increased speed of constrictions (relative to dilations) during NREM and REM, but not during WAKE. **g.** Event-triggered averages of pupil size in NREM, time-frequency activity, HR, BR, muscle tone, and the probability of occurrence of arousal and K-complexes (from top to bottom) when considering constrictions (N=19, n=47, 5382 events), dilations (N=19, n=47, 6016 events), K-complexes (only in NREM2; N=19, n=47, 4170 events), arousals (N=19, n=45, 930 events), and infra-slow oscillations (N=19, n=47, 2381 events) as triggering events (from left to right). Thick horizontal lines on line plots and dark contours on heat maps represent significant differences against baseline tested by cluster-based permutation statistics.

We next analyzed the relationship of pupil size with physiological parameters controlled by the autonomic nervous system (Fig 2c). We found that heart rate is tightly coupled to pupil size during NREM (significant positive correlation cluster from −13.1 to 0.2 s; t_cluster_=719.944, p<0.001 and significant negative correlation cluster from 2.5 to 13.3 s; t_cluster_=−513.966, p=0.002; Fig 2c and 2d). Similarly, we observed that breathing rate covaries strongly with pupil size (strongest cluster from 5.7 to 15.2 s; t_cluster_=−324.403, p=0.020). Taken together, we show that the pupillometric measures provided by the iSleep monitoring system offer an accurate proxy for brain activity and a reliable read-out of autonomic information in sleeping humans.

Whereas cross-correlation analysis provided insights into continuous interactions, pupil dynamics are marked with sequences of phasic constrictions and dilations (Fig. 2e). We therefore examined if these events indicated specific patterns of brain activity and if they were related to specific changes in other physiological markers.

We found that at the onset of pupil constrictions, EEG power decreased across all frequency bands, followed by an increase in slow-wave delta and spindle-related sigma activity (significant negative cluster from 0.1 to 24.9 s; tcluster=−37339.608, p<0.001; Fig 2g second column, second row). These changes were accompanied by decreases in heart rate (from 0.1 to 22.6 s, tcluster=−797.464, p<0.001), as well as decreases in breathing rate (from 0.1 to 24.9 s, tcluster=−982.514, p<0.001), muscle tone (from 1.8 to 17.0 s, tcluster=−359.463, p=0.002), and arousal probability (from 0.1 to 24.9 s, tcluster=−1215.356, p<0.001). The results indicate that pupil constrictions during stable, deeper sleep stages (NREM) coincide with measures of increased parasympathetic pressure. In contrast, pupil dilations coincide with a reduction in sleep depth, as signaled by the broadband increase in EEG power (from 0.1 to 24.9 s, t=19213.643, p<0.001) a decrease in delta (from 3.0 to 24.9 s, t=−4002.791, p=0.001) and sigma (from 3.7 to 21.1 s, t=−1621.286, p=0.011) activity, paralleled with transitory increases in heart rate, muscle tone, and microarousal probability (Fig. 2g).

Phasic pupil events thus reliably reflect opposing neurophysiological processes, with pupil constrictions indicating the beginning of sleep consolidation and pupil dilations reflecting moments of higher arousability.

During wakefulness, transient pupil dilations also reflect increased sympathetic inputs through projections from the locus coeruleus [1], [9], whose activity is thought to contribute to the emergence of micro-arousals during sleep [28], [30]. Elevated sympathetic activity and frequent arousals have thus been proposed to be implicated in various sleep disorders [28], [31]–[33]. We therefore investigated the relationship between transient pupil dilations, k-complexes and micro-arousal probability during NREM. Our analysis showed that K-complexes were concomitant with pupil dilations (from 0.1 to 24.9 s, t_cluster_=1896.682, p<0.001; fig 2g - third column) and indicated the begin of micro-arousals (from 0.1 to 21.7 s, t_cluster_=1352.746, p<0.001, fig 2g - fourth column). Similarly, during micro-arousals, the pupil diameter increased on average (significant cluster from 0.3 to 24.9 s; t_cluster_=969.609, p<0.001; Fig. 2g - second column). Finally, muscle tone, another marker of sympathetic activity, showed a significant increase at the time of pupil dilation (significant cluster from 1.0 to 9.4 s; t_cluster_=181.938, p<0.001; Fig 2g). Together, these findings support the idea that pupil dilation can be used as a reliable readout for sympathetic drive during NREM sleep.

Lastly, we asked if the continuous interplay between the parasympathetic and sympathetic activity also relates to pupil size at the infraslow (0.01-0.1 Hz) time scale, periodicity of which has been shown to reflect cyclic alterations in cortical and excitability [34], [35]. We found that infraslow oscillation (ISO)-triggered averages of pupil size revealed a clear rhythmicity (significant cluster of negative correlation from 2.4 to 32.9 s; t_cluster_=−1509.292, p<0.001 and significant cluster of positive correlation from 43.6 to 76.3 s; t_cluster_=1102.320, p<0.001). Other physiological measures equally showed co-waxing and waning, accompanied by the cyclic timing of arousals at pupil dilations (Fig 2g rightmost column, significant negative clusters: 0.1 to 19.8 s, t_cluster_=−15955.849, p<0.001 for power; 2.4 to 32.9 s, t_cluster_=−1509.292, p<0.001 for pupil size; 0.1 to 27.9 s, t_cluster_=−1615.989, p<0.001 for arousal probability; 1.9 to 17.8 s, t_cluster_=−534.740, p=0.005 for HR; 1.7 to 25.3 s, t_cluster_=−689.249, p=0.001 for BR and 10.1 to 19.9 s, t_cluster_=−281.454, p=0.001 for muscle tone).

Taken together, we conclude that pupil size dynamics, revealed with the iSleep system, reflect the autonomic balance throughout sleep. Continuous pupil tracking therefore provides a reliable readout of tonic parasympathetic pressure, as well as of sympathetic activity during phasic periods of increased arousal.

### Reliable prediction of sleep stages with iSleep

With the increased use of large databases, machine learning algorithms and artificial intelligence, automated sleep scoring is steadily improving [36], [37], providing the basis for accurate sleep tracking and disease monitoring. Building upon the above identified close association between pupil size and brain states, we explored the predictive capabilities iSleep to identify specific sleep stages.

We initially extracted features derived from pupil dynamics, including differentials, distributions, frequencies, and wavelets, as well as the horizontal and vertical signals of the pupil center, within 30-second windows. Using unsupervised dimensionality reduction techniques, we found that the extracted features formed clusters according to the sleep stages according to the t-distributed stochastic neighbor embedding (t-SNE) method (Fig 3b). We subsequently trained machine learning algorithms for each session by associating these features with manually-scored sleep stages, following the guidelines of the American Academy of Sleep Medicine (AASM) [30]. Using the bagging trees method [38] for training, we validated the resulting predictive values using a 5-fold cross-validation approach (N=19 participants, n=58 sessions, Fig 3a).

**Figure 3.**
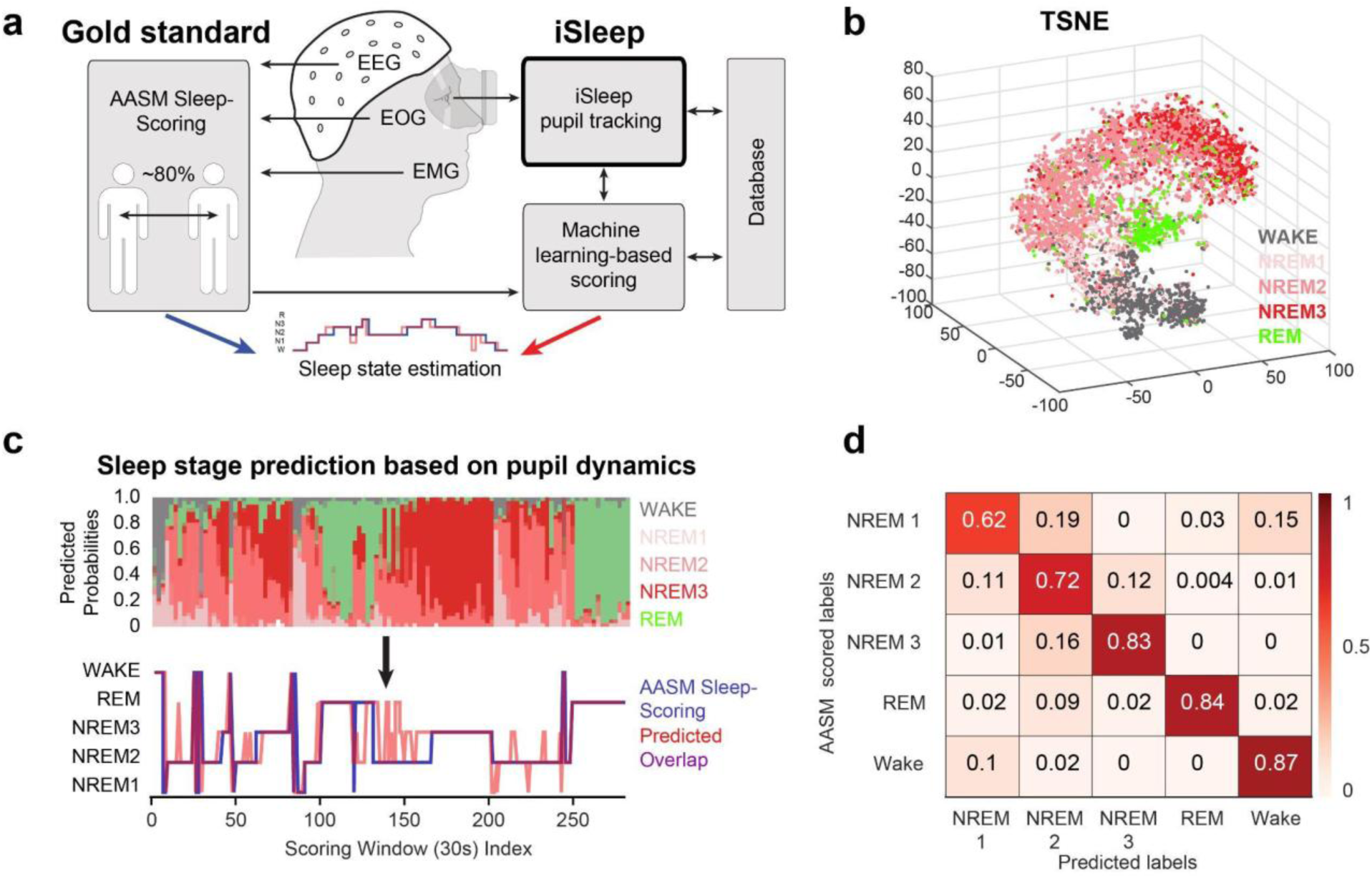
Sleep stage predictions based on iSleep data and ML algorithms. **a.** Pupil-based sleep stage detection framework and its comparison to the gold standard AASM manual scoring. **b.** Separation of the pupil-based features as a function of sleep stages displayed in t-SNE plot. **c.** Distribution of sleep stage likelihood in an example session based on pupil parameters (up) and the corresponding hypnograms for “winning” sleep stage predictions and manually scored PSG data (down). **d.** Confusion matrix for standard PSG and eye-data-based classification of sleep stages, with decoding accuracy.

Across all sleep-wake stages, prediction accuracy was 83% for NREM3, 84% REM and 87% WAKE, and slightly lower for NREM2 and NREM1 (72% and 62%, respectively)(Fig 3d, Fig 3S, Supplementary Table 3). These numbers are comparable to the expected percentage of agreement for manual scoring between two trained experts [39]. Increasing the window size from the AASM standard of 30 s to 60 s, further increased the prediction accuracy (i.e., matching with the AASM manual scoring [39]; Fig S3). Longer windows likely minimize the effects of unstable sleep periods while better mimicking AASM-based scoring decisions that require the assessment of data from neighboring temporal windows [40]. Lastly, we developed a general model by using the data from all 19 participants (Fig S3). Our results show that the generalized model is also robust with average balanced accuracy above 71%. Combining different sleep stages further improved our models, with accuracies reaching beyond 85% in agreement to the manual scoring (Fig 3d, Supplementary Table 3).

Taken together, tracking pupil size and eye movement with iSleep provides a non-invasive and automated way to detect sleep stages with accuracies comparable to the current gold standard method of manual scoring.

### Visually evoked responses support the role of pupillary constrictions as a peripheral sleep stability mechanism

Sensory stimulations have not only been used in the investigation of sleep mechanisms and maintenance, but also in diagnosis as well as treatments of various sleep and consciousness disorders [41]–[43]. Sleep stability when faced with sensory stimulations is ensured by central and peripheral mechanisms that operate during different sleep stages [44], [45]. For example, peripheral mechanisms include the closing of eyelids and reduced body motion to suppress visual and proprioceptive/tactile input. One main central mechanism encompasses thalamic gating, which “filters out” sensory information that could disturb sleep [46], [47]. Our previous experiments in mice revealed that, in conditions where the eyelids do not filter all visual information (or when the eyelids are open), pupil constrictions may act as an additional peripheral filter of visual stimuli, thereby preserving stability of deep sleep [10]. Here, we made use of the programmable visual stimulation capabilities of the iSleep goggles, to investigate whether pupil fluctuations may also subserve a protective function in human sleep.

We first employed a pharmacological approach to determine the mechanisms involved in the pupil size variations during sleep. Using the binocular monitoring feature of the iSleep goggles, we blocked the main afferents to the different pupil muscles in one eye, while using the other eye as internal control (Fig 4a). The sympathetic afferents were blocked using eye drops of the alpha-1 antagonist Dapiprazole (0.5%), whereas the parasympathetic inputs were inactivated in separate experiments using a blocker of the muscarinic system (Tropicamide, 0.5%). The constricting effect of dapiprazole was small, yet significant during sleep (mean pupil diameter change= 0.323 mm, p=0.023, student’s t-test). Parsing out the individual sleep stages revealed that the effect was visible only during WAKE and NREM1 stages and not significant otherwise. On the other hand, blocking the parasympathetic system with Tropicamide increased pupil size to its maximum across wake and all sleep stages (mean pupil diameter change: 5.940 mm, p < 0.001, student’s t-test) (Fig 4a-b). As a consequence, the correlation between pupil size of each eye was weaker for the tropicamide condition (p=0.044, student’s t-test, Fig 4b), as compared to the sympathetic blockage. The remaining fluctuations were likely due to incomplete blocking caused by the low doses used. Taken together, these results suggest that parasympathetic activity predominantly determines the constriction of pupil size during deeper, stable stages of sleep.

**Figure 4.**
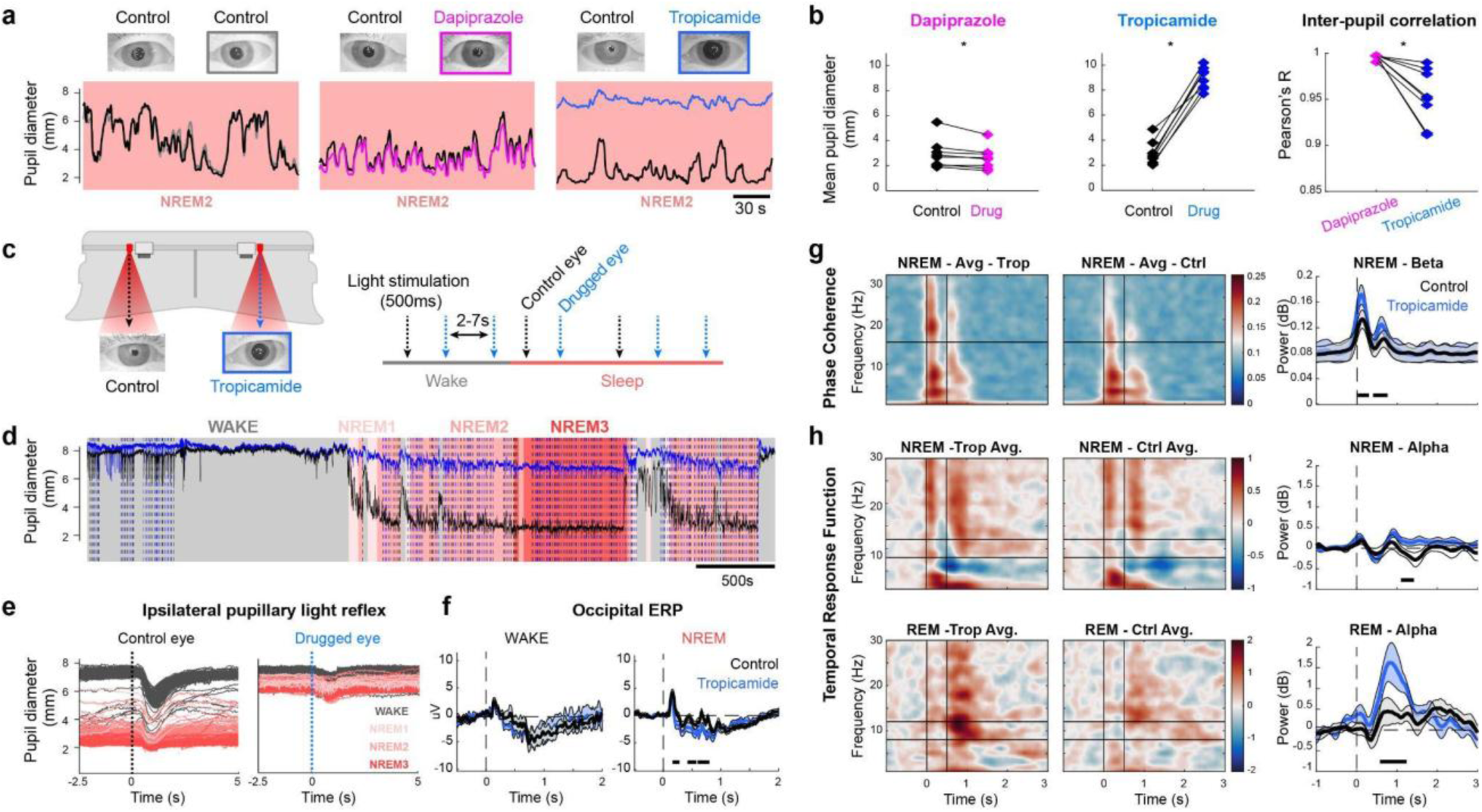
Programmable, targeted light stimulations with iSleep reveals the protective role of pupil constriction during sleep. **a.** Example traces of pupil diameter and corresponding eye video snapshots during NREM2 sleep in a baseline control where both pupils were intact (left), in dapiprazole session, where one pupil was constricted due to the sympathetic blocker (middle, pink line); a tropicamide session where one pupil is dilated due to the parasympathetic blocker (right, blue line). **b.** Mean pupil size comparison between the control and drugged eye in dapiprazole (left) or tropicamide sessions (middle). The cross correlation between control and drugged eyes in dapiprazole and tropicamide sessions (right). **c.** Light stimulation experiment setup, with monocularly targeted LEDs (left) and trial structure (right). **d.** Pupil diameter, light stimulation timings and sleep stages in the example session. **e.** Pupillary response to light stimulations in control (left) and drugged eye (right) across different sleep stages in the example session. **f.** Average of occipital ERPs time-locked to the light stimulation for the control (black line) and drugged (blue line) eyes in WAKE and NREM states. **g.** EEG beta band activity phase alignment difference on all electrodes in responses to light stimulation between drug (left) and control (middle) trials, with significant increase of beta synchronization in tropicamide trials (right). **h.** Time frequency plots of cortical responses to light stimulations across all electrodes for drug (left column) and control (middle column) trials, during NREM (top row; drug: N=6, n=18, 3396 events, control: N=6, n=18, 3326 events) and REM (bottom row; drug: N=4, n=8, 329 events, control: N=4, n=8, 316 events) sleep. Difference between the drug and control trials in the alpha band is displayed in the line plots (right column).

We next used the iSleep’s visual stimulation feature to investigate how pupil constrictions affect the processing of visual information during sleep. We temporarily dilated the pupil in one eye by applying Tropicamide eye drops, while the other intact eye was used as control (Fig 4c). Lateralized red light flashes (500 ms, 10 uW) were presented randomly to the left or the right eye every 2 to 7s to investigate the role of naturally occurring pupillary fluctuations in sleep protection. (Fig 4c,d). The stimulations began after the first occurrence of k-complexes in NREM sleep, and continued across all stages of sleep. Stimulations were halted upon the participant entering a WAKE state and subsequently resumed when NREM1 sleep was reached again. During this experiment, we simultaneously recorded EEG, EMG, ECG, EOG and iSleep pupil data.

We observed that light-evoked pupil constrictions were maintained throughout sleep for both treated and control eyes, although at significantly different amplitudes (Fig 4e). Visually evoked event related potentials (ERPs) measured by EEG showed a larger response in the dilated as compared to the control pupil during NREM (Fig 4f). These findings were subsequently confirmed by analyses of time-frequency decomposed brain responses, which revealed that stimulation of the treated eye elicited a temporally more stable transduction of visual information from the retina to the cortex (i.e., increase in intertrial phase coherence (ITPC) in the beta frequency range; from 0.02 to 0.30 s, t_cluster_=56.262, p=0.009 and from 0.40 to 0.76 s, t_cluster_=68.382, p=0.004; Fig 4g). In addition, light stimuli evoked more arousal-related activity in the alpha power, when presented to the dilated eye than in the control eye in both NREM (from 1.10 to 1.42 s, t_cluster_=45.610,p=0.015) and REM sleep (from 0.58 to 1.24 s, t_cluster_=98.807, p=0.011; Fig 4h). Taken together, these results indicate that neural responses to visual stimuli are reduced during more stable parts of sleep, when pupil size is at minimum, while neural responses increase when the pupil size is pharmacologically enlarged. Vision research has demonstrated that large pupils typically increase light influx as well as detection of peripheral stimuli, while small pupils can increase visual acuity [48], [49]. Accordingly, we suggest that small pupil size (and pupil constrictions) promote sleep stability by reducing the neural impact of external light stimulation.

### Binocular gaze tracking with iSleep reveals disconjugate eye movements in NREM and REM sleep

In wakefulness, eyes usually move in a conjugate manner during saccades, smooth pursuit, and vestibulo-ocular reflex movements. These movement categories are related to specific goal-directed behaviors such as scanning of visual scenes, following moving objects and retinal stabilization [11], [50]. Eyes also move during sleep. This is particularly pronounced during rapid eye movement (REM) sleep, hence the generally accepted nomenclature. Indeed, when scoring sleep stages, the distinction between drowsiness/NREM1 and REM sleep typically relies on the detection of EOG-based slow versus rapid eye movements, respectively. In previous studies, the relationship between eye movements and dreams or other cognitive content has been studied [51], [52]. However, these findings are based on EOG data from standard PSG recordings that provide limited information about spatially precise eye-positions. The binocular optical tracking system of the iSleep system ideally provides access to the full extent of eye-movement parameters during sleep. As for pupil size fluctuations (see Fig. 1-4 above), here we report precise eye-position measurements across distinct sleep-wake states.

Previous studies during wakefulness have already indicated that with drowsiness, the coordination between two eyes starts decreasing [53]. It has also been suggested in a primate study that during REM sleep, eye movements become disconjugate [54]. We therefore analyzed the horizontal and vertical alignments of both eyes. By leaving an error margin of 5.5% (∼1.38 mm), we first defined our boundaries of alignment as the absolute difference between the center of two eyes being within the limits.

In awake participants, the eyes appeared aligned in both horizontal and vertical coordinates more than 95% of the time, whether they were fixating objects or actively exploring visual scenes in the far distance (objects located >20m) or in a indoor setting (objects located <5m, Fig 5b,c). Strikingly, as the participants shifted into sleep, the eyes displayed “unnatural” positions, showing convergent and divergent positions/movements in both horizontal and vertical dimensions around 50% of the time (Fig 5a). Such disconjugate eye movements are usually only observed in patients with injuries at the level of the peripheral eye motor control or after brainstem damage [55]–[57]. Throughout sleep sessions, the eyes were found to be 28% in convergent and 25% in divergent positions. In NREM1, they leaned towards divergence (35% of NREM1), potentially due to eyes rolling up (known as the Bell Reflex) while in NREM2 and REM they appeared more convergent (34%, 30% of the respective stages); and in NREM3 conjugate 57% of the time (Fig 5a,c). The coordination in the Y axis remained more robust than in the X axis during sleep (Fig 5a,c). These findings suggest the mechanisms and reflexes governing conjugate eye movements when awake, are largely disrupted during extended periods of sleep. The use of binocular eye movement tracking is thus not limited to the scoring of REM sleep, but bears the potential to detect other physiologically-relevant events in all sleep stages, which have so far been hidden behind closed eyelids.

**Figure 5.**
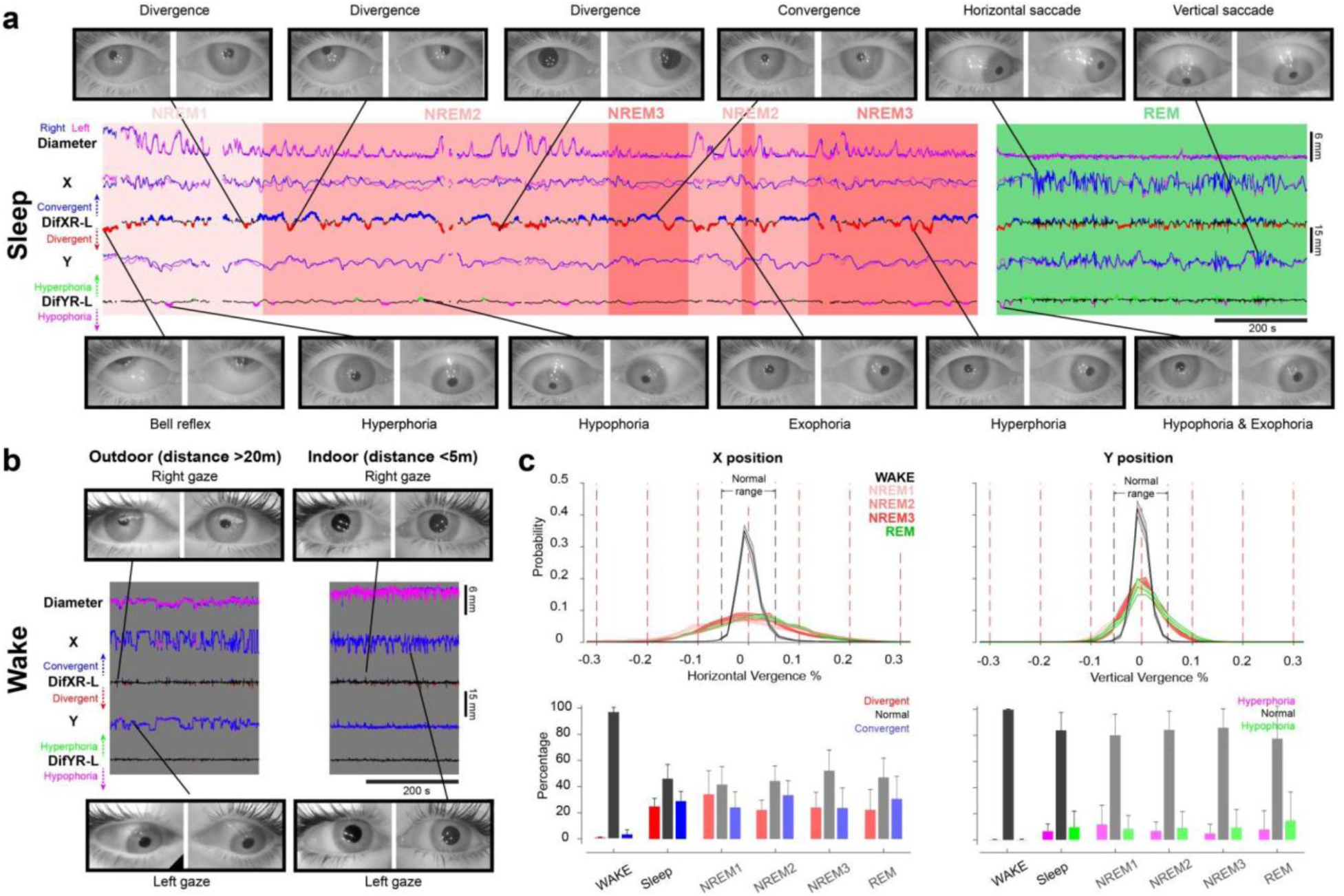
High resolution video imaging of the eyes reveals abnormal positions during sleep. **a.** Continuous traces of eye movement parameters in a typical session, with snapshots depicting divergent positions, bell reflex and saccade examples as well as other abnormal positions. Note that the right eye was taken as reference when describing abnormal positions. **b.** The eye movement traces of the same participant during wakefulness with snapshots depicting aligned eye positions. **c.** Quantification of the x (left column) and y positions (right column) of the eyes in sleep and wake sessions (N=7 participants, n=10 sessions) in comparison to sleep sessions in which both eyes were present (N=17 participants, 50 sessions).

### Clinical use of iSleep for pupil monitoring under general anesthesia

Pupil size and light induced reflexes are frequently examined in unconscious subjects in intensive care units, emergency departments and operating rooms [58]–[61]. At the beginning of most surgeries conducted under general anesthesia, anesthesiologists or nurses briefly open the patient’s eyelids to verify the anesthesia level, indicated by binocularly constricted pupils. In parallel, various physiological monitors are used to observe the patient’s health status during surgeries including heart and breathing rates, and blood oxygenation. In higher-risk surgeries these measurements are complemented with EMG, EEG and other neuromonitoring devices to help clinicians make informed intraoperative decisions [62], [63]. Despite these efforts, it remains a challenge to have reliable, automated and non-invasive handles to the depth of anesthesia or consciousness states during surgeries [64], [65].

Based on discrete and manual pupil assessment during anesthesia, it has been hypothesized that pupil size might be impacted by different drug combinations [66]–[68]. Additionally, transient observations of pupil size in relation to external stimuli under non-surgical general anesthesia has indicated that pupil dilations were induced upon noxious stimulations [69], [70]. However, how pupil size evolves during the course of surgical interventions and reacts to stimuli has so far not been investigated in a continuous manner. To address this gap, we used iSleep during neuro (anterior anterior cervical discectomy and fusion, ACDF), gynecological, visceral and plastic surgeries to track pupil dynamics under general anesthesia for up to 2 hours.

The ACDF surgeries are carried out to remove and replace damaged or degenerated discs in the neck. Known for being painful, ACDF procedures include incision and access to discs, locating and removing damaged discs, preparing bone graft fusion, inserting the fusion cage and fixing it in place, subsequently moving onto the next damaged disc if any (Fig 6b). These steps are inherently delicate as the surgeons approach intricate structures such as vital nerves and spinal cord. Hence, thorough physiological monitoring of the patient is required. We used iSleep in 3 patients during pre-surgical anesthesia setup phases as well as during 17 surgical interventions. Our system was synchronized to the surgical stimuli as indicated by the surgeon’s comments and operative camera monitors (Fig 6a-c). We observed that pupil size was dynamic throughout the surgery and numerous large dilations (0.5-3 mm) were recorded at the time of noxious surgical stimuli such as cage insertions, bone drilling, skin incisions and manipulations (Fig 6b). While the largest dilations (>1 mm) were observed at the time of skin suturing and cage insertions, we also observed small but consistent dilations (<1 mm) at the time of drilling and tissue manipulation (Fig 6b). Lastly, we compared pupil dilations in relation to the noxious stimuli, namely jaw thrust maneuver, with varying drug dosage in a pre-surgical general anesthesia setting. We observed that pupils had 4-fold increased dilations before the final additional pre-surgical opioid dosage was administered (Fig 6d). These experiments illustrate that iSleep can be easily integrated into operating room routines, allowing continuous observation of pupil size in parallel to other physiological monitoring with future potential to improve detection of anesthesia depth as well as nociceptive monitoring.

**Figure 6.**
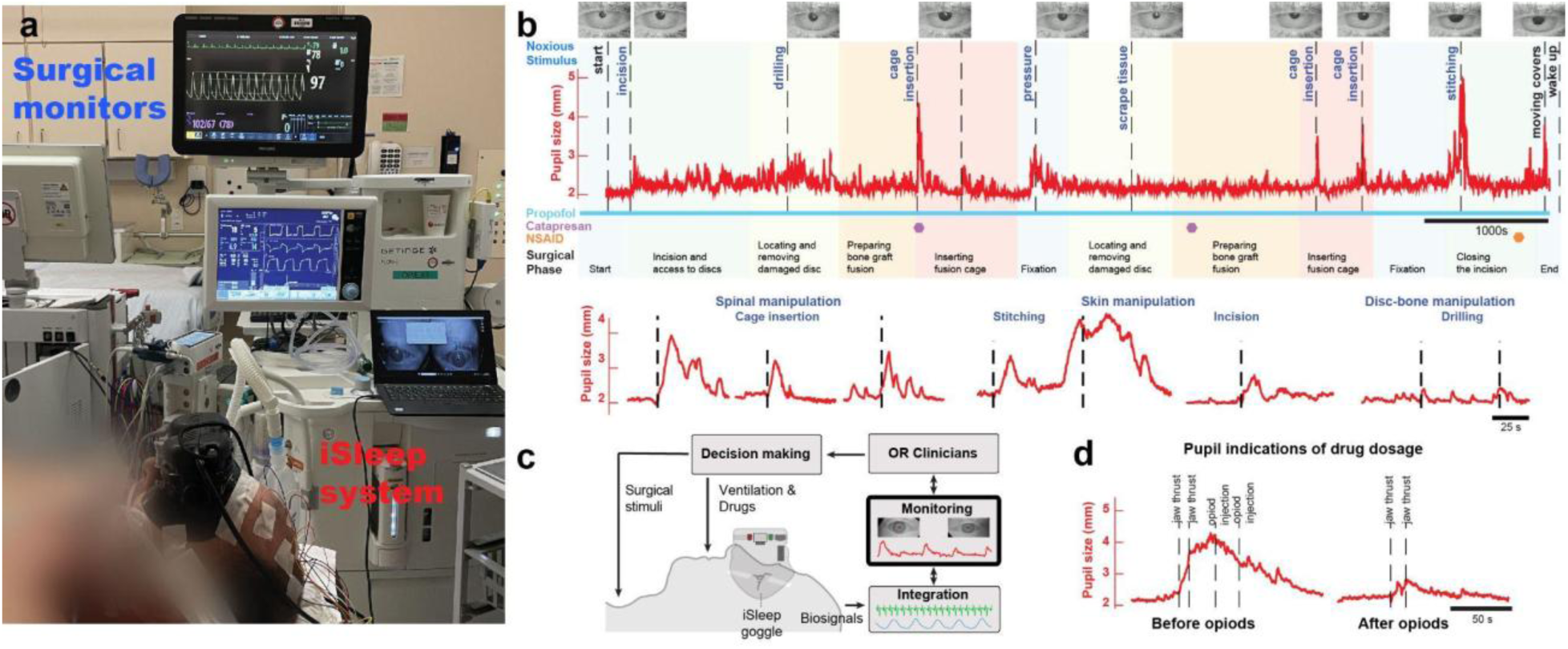
Monitoring pupil size dynamics during neurosurgery under general anesthesia with iSleep. **a.** A patient wearing the iSleep goggle after intubation prior to surgery procedure. Pupil tracking in parallel to the other standard physiological monitors in the operating room. **b**. First row: Pupil size trace, dilation examples triggered by the surgical stimulus across operational phases. Second row: Example of surgical stimuli: cage insertion (left) and skin suturing (right). **c.** Schematic representation of the utilization of iSleep in clinical settings, with the goal of helping intraoperative decision making. **d.** Comparison of pupil dilations to the same noxious stimulus with different drug dosages in the same patient.

## Discussion

The development of iSleep and its use in experiments during sleep and general anesthesia emphasize the utility of continuously monitoring binocular eye parameters: the eyes provide reliable and physiologically relevant information beyond wake periods, when the eyelids are naturally open. We showed that pupil tracking in sleeping humans with form-fitted and humidified goggles is safe, comfortable and achievable within one single session, giving continuous access to both pupil size and eye movements at high resolution. Having access to both these parameters is currently not attainable through only using EOGs, coils or ultrasound imaging [68], [71]–[73].

In this study, we conclude that pupil fluctuations are continuously regulated by the parasympathetic pathway during deeper NREM and REM stages, while during wakefulness and light sleep sympathetic input can also be inferred from the pupil size. As a result, pupil changes can predict sleep stages based on average size, whereas arousals can be predicted by the rapid, transient dilations. Supporting this view, we deduce that pupil dilations and constrictions reflect complementary mechanisms showing arousal and sleep onsets, respectively. Additionally, using high resolution IR-imaging capabilities of iSleep we described a variety of abnormal eye positions during sleep. Considering pontine substrates that are implied in binocular gaze control [74] and sleep modulation [75], these ocular observations may have physiologically relevant indications such as local arousals or absence of synchronized brain activity. These dynamics could be used to conduct further investigations linking EEG-brainstem activity in sleep research and beyond in health and disorders [13].

The current gold standard for identifying brain states in both rodent and human research is the use of EEG recordings. This approach is comprehensive and reliable, yet it can be costly and complex to implement. Complementary to EEG, other methods like monitoring heart rate, skin conductance, and breathing rate provide valuable insights, though they often respond more slowly compared to the rapid dynamics of brain states. Optical pupil tracking, utilizing image processing, emerges as a cost-effective enhancement to these techniques. Leveraging the latest open-source toolboxes, it offers a promising approach. Notably, in terms of sleep and arousal prediction, pupil size measurement has shown accuracy levels that are on par with those achieved by EEG-based scoring [74], [75]. The differences in accuracy between sleep scoring with pupil data vs. PSG may even highlight the nuances in sleep regulation that are not fully reflected in the EEG and other biosensor signals. This suggests that, when used in conjunction with traditional methods, pupil tracking can enrich the overall understanding of brain states.

iSleep provides researchers not only the means to investigate close relationships between pupil size, brain and autonomic states, but the findings presented here also confirm the role of pupils as a sleep protection mechanism and as a way to interact with the brain of a sleeping person. These findings are comparable to other experiments during which other sensory modalities were tested during sleep. In these studies the modulation of conscious processing and sleep stability was determined by cortical states and stimulus relevance [76]–[78]. Having access to open eyes during reduced consciousness and sleep, novel applications can now be designed to test perception, responsiveness and brain states, potentially becoming a game changer for novel therapeutics and diagnostics in neuroscience, psychology and medicine.

Finally, the use of iSleep in clinics and in mobile applications could enable large-scale diagnostics. Further developments of the iSleep system, using alternative methods to access the pupil through the eyelids would be beneficial to make it widely available in not only well controlled research and clinical settings but also for home use or mobile applications. The main current limitation of the system is having to keep the eyelids open, which without proper training may lead to corneal damage, eye dryness or inflammations. However, on users with confirmed healthy eyes under controlled conditions this application did not produce adverse events, even upon repeated use during sleep. Building upon this high-resolution, ground-truth data; hardware changes to achieve closed-eye pupil tracking would help eye-parameters become accessible, reliable and non-invasive biomarkers for many fields of research and medicine. Further engineering effort and adaptations will be necessary to establish such studies, yet with exponential advances in technology this might be achievable in the blink of an eye.

## Methods

### iSleep framework

The iSleep framework includes (i) form-fitted, humidified and temperature controlled goggles with real-time monitoring and stimulation capabilities, (ii) an integrated database of sleep-pupil data with synchronized biosignals and (iii) machine learning-based algorithms for brain-state monitoring based on pupil parameters. For the comfort of the user and to contain the humidity, iSleep goggle consisting of a face-fitting silicone mask were adapted from commercial diving masks (Decathlon Subea). To create space for a humidity chamber as well as to provide enough distance between the eyes and recording cameras, an internal scaffold was designed (SolidWorks), 3D printed (Ultimaker 2+ FDM printer) and mounted onto the mask frame. The eye-tracking, stimulation, temperature-humidity related electronic components were mounted on an integrated printed circuit board (PCB). The PCB included control electronics, temperature (Texas Instruments LM35) and humidity sensors (Honeywell HIH-5030/5031), six scattered 880 nm infrared light sources (6 mW placed ∼50 mm from the eyes), two 650 nm red LEDs, Pupil Labs add-on infrared eye tracking cameras (640×480 pixels, acquired at 30 Hz)[79] and a microcontroller (Seeeduino Xiao) to deliver triggers (Fig. 1a-b). The PCB then was mounted onto the internal 3D printed scaffold. To place humidifying elements inside, two 30mm x 10mm sponge holders were minted onto the internal scaffold. Finally, to protect all electronic components an external shell was 3D printed and mounted onto the internal scaffold.

For monitoring and recording the eye images in real-time, Pupil Labs Capture software was used, running on a portable computer (Lenovo x280). For monitoring temperature-humidity sensors, synchronizing datastreams and controlling light stimulations; Arduino IDE software serial monitor was used to interact with the iSleep goggle microcontroller. Based on the commands sent to the microcontrollers through the Arduino serial monitor, TTL triggers were generated by the iSleep google, which were subsequently sent to the separate PSG acquisition system (see below). In addition, infrared LEDs flashing for 1s were used to mark the acquired eye images at the time of TTL triggers to synchronize the different recording streams. Synchronized data streams were then saved in the local hard drives, which were transferred to the storage servers for preprocessing, analysis and algorithm development.

### Open eye sleep procedure

PSG preparation was conducted identically in open-eye and closed-eye naps (see below). After installation of PSG, the participants laid in the bed and their eyelids were taped open up to 10 mm by the experimenter using 15 mm x 40 mm Hypafix stretch medical tape. Three pieces were used for the upper eyelid and one for the lower eyelid. Describing the procedure to the participant and rapid application of the tapes were found helpful for the comfort of the participant and the quality of eye recordings. Two pieces of single-use viscose sponges were soaked in boiled water and were placed in the 3D printed sponge holders of the iSleep goggle. The participants then wore the goggles, were given earplugs for noise isolation and a radio-transmitting buzzer to report in case of an emergency during their nap.

All procedures were approved by the Geneva Cantonal Commission of Research Ethics (CCER) and were assigned the project ID 2019-01906.

### Experimental design

#### Recruitment process

Healthy male and female individuals ranging from 18 to 40 years old, with no history of neurological, sleep or eye problems were recruited for the study. Individuals who scored lower than 5 on the Pittsburgh Sleep Quality Inventory [80] and complied with the demographic criteria (non-smoker, no regular medication, no daily consumption of alcohol and drugs) in the demographic questionnaire were invited for a 2-hour long assessment nap. Individuals who had at least 1 h total sleep with a sleep onset <30 min, showing no signs of physiological abnormalities in the PSG data continued with the ophthalmologic assessment. Individuals with no ophthalmological disorders were invited for a 2-4 hour long open-eye nap assessment. If they were able to sleep for a minimum of 70% of the session they were enrolled for the study.

A total of 59 participants fitting the inclusion criteria (18-40 years, healthy, good sleepers, without any known eye problem) were recruited. 44 participants were confirmed to have healthy eyes and good closed-eye sleep, hence were consequently admitted for the open-eye habituation nap to evaluate their suitability for pupil tracking. 35 participants slept with their eyes open without any disturbance and for longer than one hour, and thus were admitted to the study, resulting in a 80% retention rate of iSleep. The reasons for exclusion from the study included fragmented sleep, too frequent blinks or particular anatomy of the frontal bone. The data from the first 16 participants was used to set up and refine the procedures. It contained tests with different camera setups and various PSG arrangements. All data from the subsequent 19 participants were included.

#### Ophthalmic examinations

Before and after the open-eye naps, participants went through the following exams to assess their eye health. The list of parameters that were used to determine the health of the participant’s eyes were:

Intraocular pressure within the range of 10-21 mmHg, visual acuity of 10/10^th^, and normalcy of the eyelids, conjunctiva, cornea, anterior chamber and posterior segment. The latter was evaluated using Schirmer’s test, with a minimum measurement of 15 mm after 5 min. The anterior chamber was deemed within normal limits, with the absence of hyphema, nor signs of inflammation as cellular tyndall, fibrin, or hypopyon. Fundus was evaluated through fundus photography in search for abnormalities such as signs of inflammation or bleeding within the vitreous and retina. Exclusion criteria encompassed participants more than 5 diopters of correction, inflammation, such as conjunctivitis, keratitis or uveitis, endophthalmitis, vitreous hemorrhage, choroidal or retinal detachment, retinal hemorrhage, or a history of previous eye surgery. Pupil size and reactivity symmetry, eye motility in all quadrants, and Bell’s reflex (up and outwards movement of the eye when attempting to close it) were also assessed for normality. This methodological approach ensured a thorough and systematic evaluation of ocular health, encompassing diverse facets from intraocular pressure to external manifestations and functional indicators. Eligible participants were accepted to the study. All ophthalmological assessment took place at the University of Geneva Hospital Department of Ophthalmology.

#### Closed-eye naps

Polysomnographic data was collected with 16 channel or 32 channel recording systems (V-amp 16, BrainAmp 32 Brain Products, respectively). The PSG data included EEG (F3, F4, Cz, Pz, O1, O2, A1, A2, Ground between F1 & F2 and Reference between Fz & Cz according to the 10-20 system), EMG (2 electrodes on the chin), EOG (2 channels, upper left and lower right eye) and ECG (2 channels, diagonally placed between the right collar bone and below the left ribcage or a single electrode below the left shoulder blade). During the procedure, participants slept in a sound isolated, dark room. All sleeping sessions took place at the University of Geneva Brain and Behavior Laboratory sleep rooms. They started between 09:00 and 16:00 and lasted approximately 2 hours.

#### Open-eye naps

Participants’ eyelids were gently opened with medical tape up to ∼10 mm as described in the open-eye procedure above. Participants wore the iSleep goggles and eye images were recorded. PSG data was simultaneously collected, as described above. All sleeping sessions started between 09:00 and 16:00 and lasted between 2-4 hours. The open-eye sleeping sessions were scheduled with 48 hours timing in between for each participant.

#### Pharmacological blockage experiment without light stimulations

Participants were prepared for an open-eye nap session as described above. Thirty minutes before the session, one of the eyes was instilled either with Tropicamide (Théa Tropicamide 0.5% SDU Faure, Tropicamidum 5 mg/ml) or Dapiprazole (OmniVision Glamidolo 0.5%) while the other eye was instilled with humidifying eye drops (Théa Lacrycon Acidum hyaluronicum 0.14 mg/ml). The recording continued as a regular open-eye nap while all PSG data was collected in parallel to pupil tracking.

#### Visual stimulation during sleep

Participants were prepared for an open-eye nap session with Tropicamide instillation in one eye as described above. When the participant entered the NREM1 phase of sleep, they were presented with brief light pulses (500 ms, 650 nm, 10 μW), randomly presented to the left or the right eye at intervals ranging from 2 to 7 s. If the participants woke up, flashes were stopped and were continued again with the start of NREM1 sleep, after the first K-complex.

#### Wake eye-tracking

The original iSleep goggle was modified for use in wakefulness. The camera and IR light positioning was left intact to have a comparable recording angle of the eyes. To give visual access to the participants, the external shell of the goggle was removed and PCB was adjusted to provide viewing gaps at the front of the goggle. The opening of the lateral ends of the upper eyelids were reinforced with a single, soft, elastic tape on each side to reveal the lateral commissures, which facilitated the use of the same DLC analysis pipeline in all recordings. The participants completed (i) visual exploration of decorative items (furniture, lamps, bottles, paintings etc.) within a 5 m x 6 m space, (ii) visually followed a target stick or (iii) looked outside from the windows to explore the exterior scene. The recordings lasted between 10-20 mins.

### Sleep scoring

Classification of sleep into Wake, NREM1, NREM2, NREM3 and REM phases and the scoring of micro-arousals was done according to the AASM manual [40] by an experienced sleep scorer.

### Pupil tracking

DeepLabCut [81] toolbox was used to detect pupil diameter and pupil centroid from the acquired eye images (680×400 pixels). To train the DeepLabCut network 20 frames from 10 subjects were chosen and 15 points (4 on the circumference, 1 in the center, 10 around the eyelids) were hand annotated to create an eye skeleton model. Eye size was estimated as the distance between the medial and the lateral canthi, eye center was estimated as halfway between the two canthi, the rotation angle ‘a’ was defined by the sinus of the line that connects the medial and lateral canthi. The eye frames were then subsequently registered by rotating the images a degrees and centering the frames to the eye center. The extracted eye values were then normalized by assuming the eye size to be identical and equal to 1 unit (∼25 mm). Pupil diameter was estimated as the distance between lateral and medial points detected on the pupil. Convergent eye positions were defined as instances where absolute distance between the pupil centers were within less than %5.5 (∼1.375 mm) proximity to each other. Divergent periods were defined as instances where the absolute distance between the pupil centers were within more than %5.5 units away from each other. Remaining eye positions were considered as aligned.

### Detection of large pupil dilations and constrictions

To detect large pupil dilations and constrictions, the pupil-size signal was filtered between 0.01 and 0.2 Hz with a 2^nd^ order butterworth filter and the instantaneous derivative of the signal was calculated. Subsequently, the signal was z-scored. Any part of the signal crossing the threshold of 0.25 or −0.25 of the z-scored signal was considered as dilation or a constriction, respectively The first point and the last point passing the threshold were marked as the beginning and the end of the change in pupil size, respectively. We rejected any dilation with a peak below 1 and any constriction with a trough above −1 and we rejected any dilation or constriction shorter than 0.5 s. Event-related potentials were locked on the beginning of dilations and constrictions.

### EEG analysis

#### EEG preprocessing

Cortical electrodes were referenced to the contralateral mastoid electrode. The EEG signal was bandpass filtered between 0.1 and 40 Hz with a 2^nd^ order butterworth filter. Artifacts in EEG signals were visually identified and removed.

#### Time-frequency power and phase clustering analyses

EEG signals of whole-nap recordings were decomposed in time and frequency using continuous wavelet transform. We used wavelets with frequency increasing from 1 to 30 Hz by steps of 1 Hz and with a corresponding Full-Width at Half-Maximum decreasing linearly from 1 to 0.33 s (corresponding to a linear progression from 1 to 10 cycles). Then, the power of the time-frequency decomposition was extracted as the modulus of the complex number power 2 and was normalized by applying a tenth logarithm to the numbers and multiplying it by 10 (i.e. decibel transformation). For ITPC analyses, the phase of the complex numbers was extracted with the built-in *angle* function of matlab.

#### Detection of K-complexes

Detection of K-complexes was performed on the average signal of Fz and F3 re-referenced to the mastoids with a well-validated algorithm [82]. In brief, this algorithm consisted of filtering the signal between 0.5 and 4 Hz (1^st^ order butterworth filter), finding zero-crossing points and defining K-complexes as a descending zero-crossing point followed by an ascending zero-crossing point in more than 0.25 s and less than 1 s. To ensure the detection of slow-waves with large down-states, typical of K-complexes, we excluded all slow-waves with a negative peak above −50 *μ*V. To extract a continuous index of K-complex probability, a binary signal was computed where 1 s marked the detected starts of K-complexes and all other time points were marked as 0 s. Then, this signal was smoothed with a gaussian window (σ=1 s).

#### Detection of infra-slow oscillations

Infraslow-oscillations were detected by averaging normalized (through decibel transformation) power within the sigma frequency band (12-15 Hz) of the time-frequency decomposed signal of the entire recording [35]. We then filtered the sigma power between 0.01 and 0.025 Hz with a 2^nd^ order butterworth filter and extracted the phase of the Hilbert transform of the signal with the *angle* function of matlab. We defined the start of the ascending phase of the infraslow-oscillation as the zero-crossing point of the phase.

### Heart rate analysis

Bipolar ECG electrodes were subtracted from each other and the subtracted signal was filtered between 1-150 Hz. To detect heartbeat based on the peak of R waves, we used an algorithm resilient to noise and signal polarity detailed here [83]. Heart rate was calculated as beats per minute by computing the instantaneous difference between consecutive heartbeats. The signal was up-sampled to 500 Hz with linear interpolation and was smoothed using a moving average of 10 s.

### Breathing rate analysis

Individual breaths were detected using the BreathMetrics matlab toolbox with default parameters [84]. Breathing rate was calculated as breaths per minute by computing the instantaneous difference between consecutive breaths. Breathing rate was up-sampled to 500 Hz with linear interpolation and was smoothed using a moving average of 4 s.

### Muscle tone analysis

Bipolar EMG electrodes were placed according to standards of PSGrecordings [40]. To extract the muscle tone, the EMG signal was filtered between 30 and 100 Hz with a 2^nd^ order butterworth filter and the amplitude of muscle tone was computed as the modulus of the Hilbert transform of the filtered EMG signal. To reduce the influence of abrupt changes, the muscle tone signal was smoothed using a moving average of 4 s.

### Cross-correlation of brain and physiological signals with pupil size across vigilance states

To compute cross-correlations, the pupil size signal of a single eye was segmented in periods of 60 s. Periods containing different sleep stages were discarded for this analysis to assure interpretability. Each 60 s-period was correlated with successively time-shifted 60 s-periods of average time-frequency power across P3, Pz and P4 channels using pearson’s R. The same procedure was applied with physiological signals such as heart- and breathing rate or muscle tone. All resulting cross-correlation patterns from the same session and the same sleep stage were averaged together. Statistics were computed on grand averages across sessions.

### Event-related responses to physiological events

Physiological event onsets were defined as dilation or constriction starts (see above), K-complex start (see above), micro-arousal start (as defined visually by the expert sleep scorer) and starts of the rising phase of infraslow-oscillations (see above). Signals of interest (pupil size, average time-frequency power across P3, Pz and P4, heart-rate, breathing rate, muscle tone, arousal probability and K-complex probability) were segmented in epochs going from −10 to +40 s relative to physiological events onset excepted for infraslow-oscillations starts for which they were going from −40 to +80 s relative to physiological events onset. Individual epochs were corrected by removing the average baseline value between −5 to +0 s relative to physiological event onset (and from −40 to 0 s for infraslow-oscillations) except for micro-arousal and K-complex probability. All resulting epochs from the same session and the same sleep stage were averaged together. Statistics were computed on grand averages across sessions.

### Event-related potentials to lights

Time-frequency power on each electrode was segmented in epochs going from −1 to +3 s relative to light stimulation onset. Then, the epochs were baseline-corrected by removing the average value between −1 and 0 s relative to stimulus onset. All resulting epochs from the same session and the same sleep stage were averaged together. To compute ITPC, the following formula was applied:

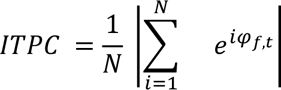

With *φ_f,t_* the phase at frequency *f* and time *t* and *i* the trial. Statistics were computed on grand averages across sessions.

### Statistical testing on continuous signals

To correct for multiple comparisons, statistical differences were computed through cluster-based permutation analyses. Clusters were defined with paired t-tests or t-tests compared to baselines values with an alpha threshold of α=0.05. N=5000 permutations were randomized to compute the cluster distribution and the cluster p-value was computed two-tailed. Clusters with a p-value below 0.05 were considered significant.

### Machine learning procedures

The streams of PSG and pupil data were preprocessed (referencing, filtering), annotated (sleep scoring, artifact rejections) and synchronized as described in previous sections. The scoring, pupil size and pupil center data streams were then epoched into 30 or 60 s windows. From the pupil data, five distinct types of features were extracted, totaling 196 features. These encompassed correlations (cross correlations of each two signal), differentials (first and second-order differentials of each signal), distribution (statistical features such as average, variance, median, skewness, kurtosis, magnitude count, and others), frequency (including maximum, minimum, and median), and frequency bands variability. Then, the feature matrix was normalized for each epoch. This was achieved by subtracting the average and then normalizing by the standard deviation of that particular epoch. Various machine learning models were evaluated using the Matlab Classification Learner Toolbox version 12.1 (R2021a). These models included decision trees, support vector machines (SVMs), k-nearest neighbors (KNNs), and neural networks (MLP). Following thorough testing, the Bagged Trees method emerged as the optimal choice. Due to the inherent imbalance in the dataset, attributed to varying proportions of sleep stages, we employed the BalancedBaggingClassifier from the imbalanced-learn library in Python version 3.7.0, utilizing a decision tree as the base estimator. The model was subsequently trained and validated on the feature matrix and corresponding scoring labels using a 5-fold cross-validation approach.

*Features list:* wavelet, bins from histogram (number of bins = 20), correlation between each two signals, maximum freq, average freq, median freq, min of signal, max of signal, median of signal, variance, std, normalized second diff, second diff, normalized first diff, first diff, kurtosis, skewness, distribution width, min to max velocity.

### Surgical recording procedures

Neurosurgery patients, who were scheduled for elective ACDF surgery and fit all of the inclusion criteria and none of the exclusion criteria, were approached by LJ or CB for a pupil tracking session under general anesthesia during their surgery. Other patients were approached by FS and LV. After providing informed consent, their eye health was confirmed by an ophthalmic exam (except for one patient, see below). General anesthesia for surgery was induced and monitored by the anesthesiologist based on the institutional standards. Anesthesia regimen usually included propofol, sevoflurane, fentanyl and rocuronium. Following administration of anesthetic agents, patients were intubated and prepared by the anesthesiology as well as neuromonitoring teams for surgery. Preparation procedures included, insertion of subdermal EMGs, EOGs, EEGs and other cutaneous electrodes for measuring heart rate, blood pressure and electrodermal activity, insertion of a urinary catheter, positioning and fixing the body of the patient in the supine position. The recording of pupil size lasted between 30-120 min.

After recording 42 subjects without any aversive incident, the entry eye exam was removed for all subjects of the study, including surgical patients, in an interim version of the ethics protocol. Within that period, one ACDF patient who had a history of eye dryness, as well as lasik surgery unknowingly got recruited in the study (post-hoc discovery of the patient background). The pupil recording under general anesthesia of this patient was registered as the only serious event of the study. It resulted in bilateral dry eyes and ocular surface inflammation. The patient was treated with Vitamin A (Bausch & Lomb 5g Retinoli Palmitas), Lacryvisc (Alcon Eye gel), Lacrycon and Dexafree (1 mg/ml Dexamethasone phosphate) over the period of 2 months, at which time point clinical symptoms were fully resolved. After this event, obligatory eye exams were reintroduced as a mandatory entry condition. In the grand scale of the study with 50 total participants and 305 sessions, no other adverse events were recorded.

## Author contributions

OY, LB, MK, CB, LV, GT, SS, and DH conceptualized the study. OY, SS, GT and DH funded the study. OY, YG, QT, SP developed hardware. OY, YG, AN, QT, CB, LJ acquired data. OY, GL, LB, AA, YG, QT, AN analyzed data. MK, AM, DL, LL, GN, SVD, GEB, GT conducted ophthalmic exams. KS, LV and GT supervised clinical recordings. OY, GL, LB, AA, AN, MK, SS and DH wrote the manuscript.

## Acknowledgements

This research was funded by the BRIDGE programme of the Swiss National Science Foundation and Innosuisse (Swiss Innovation Agency). We would like to thank the University of Geneva Brain and Behavior Laboratory (BBL) for accommodating the sleep experiments, for their technical support and organizational help. We thank all BBL personnel, especially Bruno Bonet, Sylvain Delplanque, Frederic Grouillier and Rémi Tyrand for their help with the EEG equipment and discussions. We thank Dominica de Thomas Wagner, Gregorio Galiñanes, Mario Prsa for the insightful comments and corrections of the manuscript. We would like to express our gratitude to Marco Ruedi, Elise Baud, and Callum Brindley for their guidance, discussions, and for facilitating contacts with the HUG clinicians.

## Notes

### Competing Interest Statement

The authors have declared no competing interest.

