## Supplemental figures for "iSleep: Continuous, binocular pupil tracking in sleep and reduced consciousness for physiological monitoring, predictions and interventions"

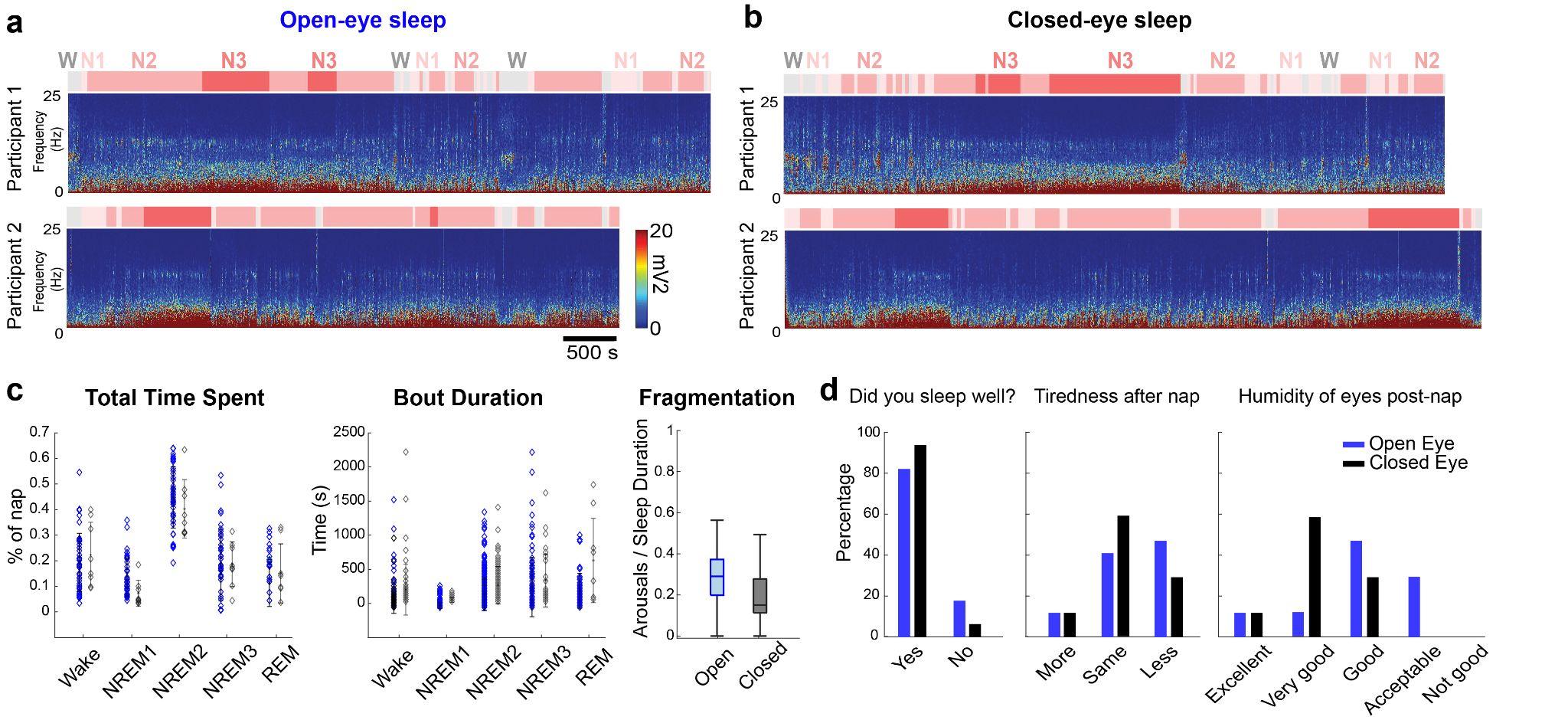

**Supplementary Figure 1. Sleep characteristics of open- and closed-eye sleep.**

**a.** Hypnograms and time-frequency plots of EEG power during open-eye naps from 2 different participants. **b.** Hypnograms and time-frequency plots of EEG power during closed-eye naps from the same 2 participants. **c.** Percentage of time spent across wake and sleep stages, stage bout durations and fragmentation quantification (in minutes) for open-eye sleep (N=19 participants, 58 sessions) and closed-eye sleep (N=17 participants, 19 sessions). **d.** Responses to post-nap questionnaires from 17 participants.

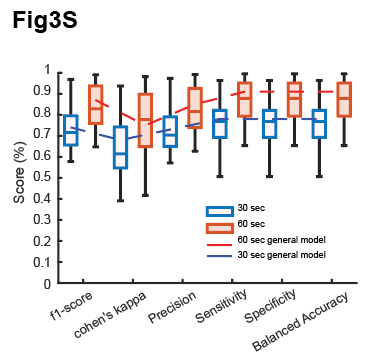

**Supplementary Figure 3. Comparison of general and personalized models with 30 vs. 60s window-based decision making.**

Average of prediction accuracies across individual sessions, calculated with models using 30s (blue) or 60s (orange) training windows. The dotted overlapping lines display the prediction accuracies for the respective generalized models. Overall, using longer time windows appears to better match AASM manual scoring, which plausibly relates to criteria involving temporal dependencies across 30s time-windows in the AASM scoring decisions.

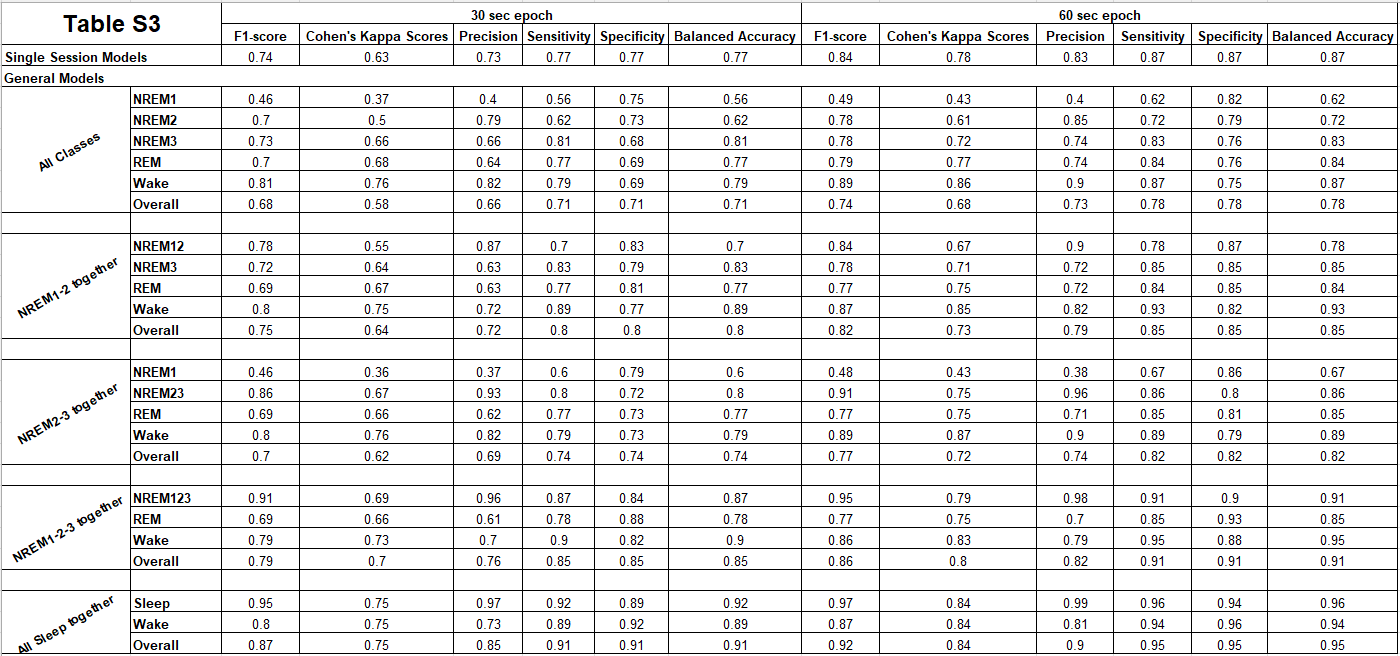

**Supplementary Table 3: Evaluation metrics for averaged single session models and the general model in sleep stage prediction with varied epoch window sizes.**

The table presents the evaluation metrics, including F1 score, Cohen's Kappa, precision, sensitivity, specificity, and balanced accuracy, for both the average of single session models and the general model (where all data is combined) predicting sleep stages based on eye tracking data. Additionally, metrics for the general model, where classes are combined, are included. The assessment covers both 30-second and 60-second window sizes, offering insights into the model effectiveness under different temporal resolutions and addressing the challenges posed by a highly imbalanced dataset.
